# Gene-Gene Interaction Detection with Deep Learning

**DOI:** 10.1101/2021.03.12.435063

**Authors:** Tianyu Cui, Khaoula El Mekkaoui, Jaakko Reinvall, Aki S. Havulinna, Pekka Marttinen, Samuel Kaski

## Abstract

We do not know the extent to which genetic interactions affect the observed phenotype in diseases, because the current interaction detection approaches are limited: they only consider interactions between the top SNPs of each gene, and only simple forms of interaction. We introduce methods for increasing the statistical power of interaction detection by taking into account all SNPs and *complex interactions* between them, beyond only the currently considered multiplicative relationships. In brief, the relation between SNPs and a phenotype is captured by a gene interaction neural network (NN), and the interactions are quantified by the Shapley score between hidden nodes, which are gene representations that optimally combine information from all SNPs in the gene. Additionally, we design a new permutation procedure tailored for NNs to assess the significance of interactions. The new approach outperformed existing alternatives on simulated datasets, and in a cholesterol study on the UK Biobank it detected six interactions which replicated on an independent FINRISK dataset, four of them novel findings.

## 1 Introduction

Genome-wide Association Studies (GWAS) have successfully identified risk alleles for complex diseases by associating single nucleotide polymorphisms (SNP) with disease-related phenotypes. Despite this, GWASs usually fail to capture statistical epistasis, i.e., interactions between genes, which has been recognized as fundamentally important to understanding the structure and function of genetic pathways. Mapping interactions between genes holds the promise of a breakthrough in revealing biological mechanisms of diseases, assess individuals’ disease risk factors, and developing treatment strategies for precision medicine^1^.

Current scalable statistical approaches lack power to reveal all gene-gene interactions, because they make restrictive assumptions about how genes are represented, and about the form of the interactions. Popular methods represent interactions between genes by interactions between the top SNPs^2^ (the SNP most correlated with the phenotype) of the corresponding genes; this ignores the effects of other SNPs within the same gene. Some scalable approaches have been proposed to summarize the multi-dimensional SNPs into a one-dimensional representation by unsupervised dimensionality reduction methods, e.g., PCA^3^, but such learned representations neglect information about the phenotype. Moreover, the interactions are modelled with parametric models, and simplifying assumptions are made when choosing the forms of the models. Linear regression with multiplicative interactions^2,4^ is a common choice. Methods based on such simplifications will fail when the actual interactions are more complex. Although more flexible machine learning algorithms, such as boosting trees^5^, have been proposed to model complex interactions between SNPs, it is still an open question how can we best combine gene representation learning and modelling of the interactions into an end-to-end model to increase the statistical power.

Deep neural networks (NNs), have been successfully applied to numerous tasks in the biomedical domain, such as predicting protein structures^6^ and promoters^7^, with large amounts of data. With a properly designed architecture, a NN learns multilevel representations by composing simple but non-linear functions that transform low-level features into abstract high-level representations, allowing the model to automatically discover optimal feature representations and approximate non-linear relations between independent and dependent variables^8^. Although NNs are effective in various prediction tasks, their black-box nature limits their usage in applications requiring model interpretability, such as GWAS. With recent progress in interpretable neural networks, interaction effects between features can be obtained by calculating well-principled interaction scores, such as the input Hessians^9^ and Shapley interaction scores^10^, which provide a way to estimate gene-gene interactions from NNs trained on GWAS datasets.

Assessing the significance of an interaction is equally important to estimating the interaction itself. As the null distribution of the interaction score is often analytically intractable for neural networks, permutation tests^11^ are used as a default solution. Such tests have been widely applied when detecting interactions with linear regression^12^. Previous approaches either 1) permute the dependent variable directly^13^ or 2) permute residuals after subtracting the null hypothesis (i.e., a linear regression without interactions) from the dependent variable^14^, and use this as the regression target when constructing the null distribution for an interaction. Notably, both of these approaches remove the main effect from the regression target. This is appropriate in linear regression where including the main effects will not change the test statistics of interactions^14^. However, unlike in linear regression, in neural networks the main and interaction effects are entangled in the high-dimensional nonlinear representations. If the main effects were excluded during permutation, NNs would learn representations that are very different from those learned from the original dataset, where main effects are included. Alternatively, they might even miss the signal altogether, which would lead to highly biased null distributions for the interaction effect.

In this paper, we present a deep learning framework to detect gene-gene interactions for a given phenotype, as well as a novel permutation procedure to accurately access the significance of the interactions. Specifically, we design a new neural network architecture with structured sparsity that learns gene representations from all SNPs of a gene as hidden nodes in a shallow layer, and then learns complex relations between the genes and phenotypes in deeper layers (Figure 1a). Interactions between genes are then estimated by calculating Shapley interaction scores between the hidden nodes that represent the genes. Moreover, we present a novel permutation test for interactions in NNs, in which the target is the sum of the permuted interaction effect (residual) and the main effect estimated by a main effect NN (Figures 1b and 1c). This helps NNs to learn gene representations in the permuted datasets that are similar to those in the original data. We employ a Bayesian approach to incorporate a prior distribution over the NN parameters to improve the accuracy and avoid overfitting. We demonstrate with simulations that the new permutation test outperforms the existing alternatives for NNs, and that the proposed deep learning framework has an increased power to detect complex interactions compared to existing methods. We then show that NNs can find from UK Biobank datasets^15^ novel significant interactions that current approaches may ignore, and we solidify our findings by replicating them in an independent FINRISK dataset^16^.

**Figure 1.**
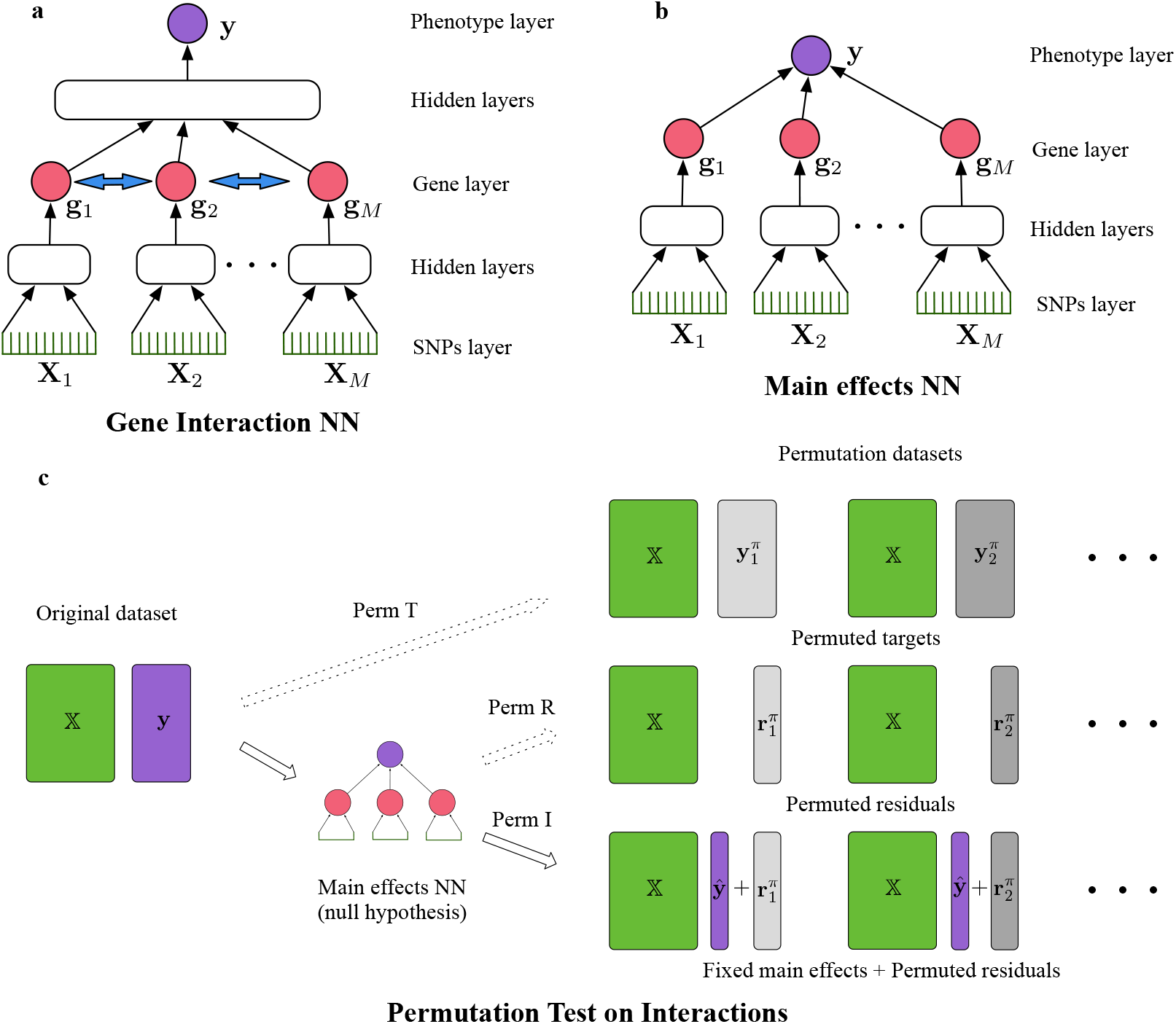
**(a)**: Gene interaction neural network to learn interactions between genes, which are estimated between nodes (blue arrows) in the gene layer. **(b)**: Main effect neural network (NN) to learn the main effects between genes and used to generate permuted datasets. **(c)**: an overview of the proposed framework. Compared with existing permutation tests (Perm T and Perm R), our new proposed permutation procedure (Perm I) includes the estimated main effects of genes from the main effect NN. Different grayscales represent different permutations.

## 2 Results

### 2.1 Novel Methods

#### Gene interaction neural network

A NN is a function mapping inputs to outputs, by composing simple but non-linear functions. Each hidden layer of the NN represents one composition. A *D*-layer NN 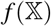 with input 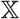 is defined as 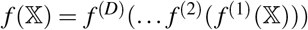, where *f*^(*i*)^(·) represents the ith layer. In a GWAS, the input 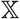 comprises SNPs and the output 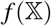 consists of the predicted phenotype for an individual. The architecture of a NN reflects the inductive biases of the model, i.e., preferences to certain kinds of functions^17^, and thus affects the learned representations. To learn representations of genes from SNPs, we propose a structured sparse NN architecture, in which SNPs from the same gene are fully connected already in lower layers, but SNPs from different genes are not connected until after a special hidden layer, here referred to as the ‘gene layer’, where each hidden node represents a single gene. After the gene layer all nodes become fully connected and finally predict the phenotype **y**, as shown in Figure 1a. The architecture reflects the domain knowledge^18^, whereby the effect of SNPs on a phenotype is assumed to follow a two-stage procedure: 1. SNPs within a gene affect how the gene expresses; 2. The combined expression of multiple genes affects the phenotype, and gene-gene interactions are estimated from the second stage.

#### Gene-gene interaction score

Shapley interaction score^10^ is a well-axiomatized measure of interaction between features that can be applied to any black-box model. To apply it to NNs, we denote the set of all input features by *F*, one feature *i* ∈ *F*, and a feature set *S* ⊆ *F*. The interaction effect between features *i* and *j*, given a set of features *S* are presented, of a NN *f* at data point *X_k_* is given by 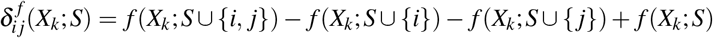, where *f*(*X_k_*; *S*) is the prediction at *X_k_* when only features in S are used and other features are replaced by their corresponding baseline values (e.g., mean)^19^. The Shapley interaction score is the expectation of 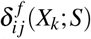 over *X_k_* and feature sets *S* sampled uniformly at random from *F*. We define the gene-gene interaction score as the Shapley interaction score between hidden nodes in the gene layer, i.e., we consider the learned gene representations as the input features of the Shapley score.

#### Permutation test for interactions

The null hypothesis here is that genes only have main effects on the phenotype and no interactions. This we formalize as the ‘main effects NN’ (Figure 1b) where only a linear layer is applied after the gene layer. To construct a permuted dataset, we first train the main effects NN on the original dataset and then permute the residual. The dependent variable in the permuted dataset is defined as the sum of the predicted main effect and permuted residual, while the independent variable is the same as in the original dataset. We also incorporate the Max-T method^20^ (i.e., collect the maximum interaction scores from permutations to construct an empirical null distribution) and false discovery rate (FDR) control^21^ into the permutation procedure for multiple testing correction. We refer the reader to the Methods 4 section for all implementation details.

### 2.2 Simulated datasets

We carry out a simulation study to validate that 1. the new permutation method outperforms existing approaches for estimating the significance of interaction, and 2. the proposed neural networks have an increased power to detect complex interactions.

#### Setting

As the genotypes, we select genes associated with the cholesterol phenotype from the UK Biobank^15^ (see Methods), from which we select all SNPs from 10 genes after basic quality control. We simulate the phenotype **y** by first simulating the expression **g**_*i*_ of each gene *i* according to a linear combination of all SNPs in the gene (Eq. 1a), and then generate phenotypes with main and interaction effects (Eq. 1b), as follows:

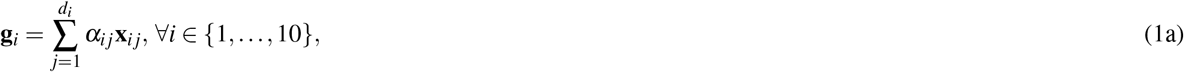

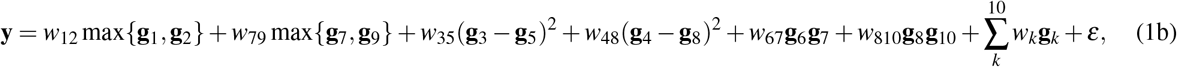

where **x**_*ij*_ is the *j*th SNP of gene *i*. This includes six interactions of three types: multiplicative, **g**_*i*_ × **g**_*j*_, maximum, max{**g**_*i*_, **g**_*j*_}, and (squared) difference, (**g**_*i*_ – **g**_*j*_)^2^. We set the weights of interactions *w_ij_* and main effects *w_k_* such that the main effect represents 80% of the signal variance, and we tune the variance of noise *ε* to adjust the signal-to-noise ratio (S/N) of the data. We consider 5 different S/N:{1.0, 0.5, 0.15, 0.10, 0.07}, which cover realistic GWAS cases, e.g., the S/N of BMI^22^ is larger than 1 and HDL^23^ has a S/N ≈0.1. We consider three different data sizes: {40*k*, 80*k*, 120*k*}, of which 70% are used for training and 30% for testing. The hyper-parameters of the NNs are chosen via 5-fold cross-validation on the training set, and the gene-gene interaction scores are computed on the test set. We compare the NN framework with 6 baselines, which are composed by 2 ways of representing genes (top-SNP and PCA) and 3 ways of modeling interactions between gene representations i.e., linear regression (LR), Lasso with multiplicative interaction terms, and boosting tree (XGBoost). We compare the new permutation methods, i.e., using permuted interactions (Perm I), with 2 existing approaches: using permuted target (Perm T) and permuted residual (Perm R) (Figure 2). We estimate the ROC curve of each algorithm on 20 datasets for each experimental setting (Figure 3 and Supplementary Figure 3), estimated by varying the p-value threshold. The p-values are obtained by the t-test in LR, and by the best permutation method for the other models.

**Figure 2.**
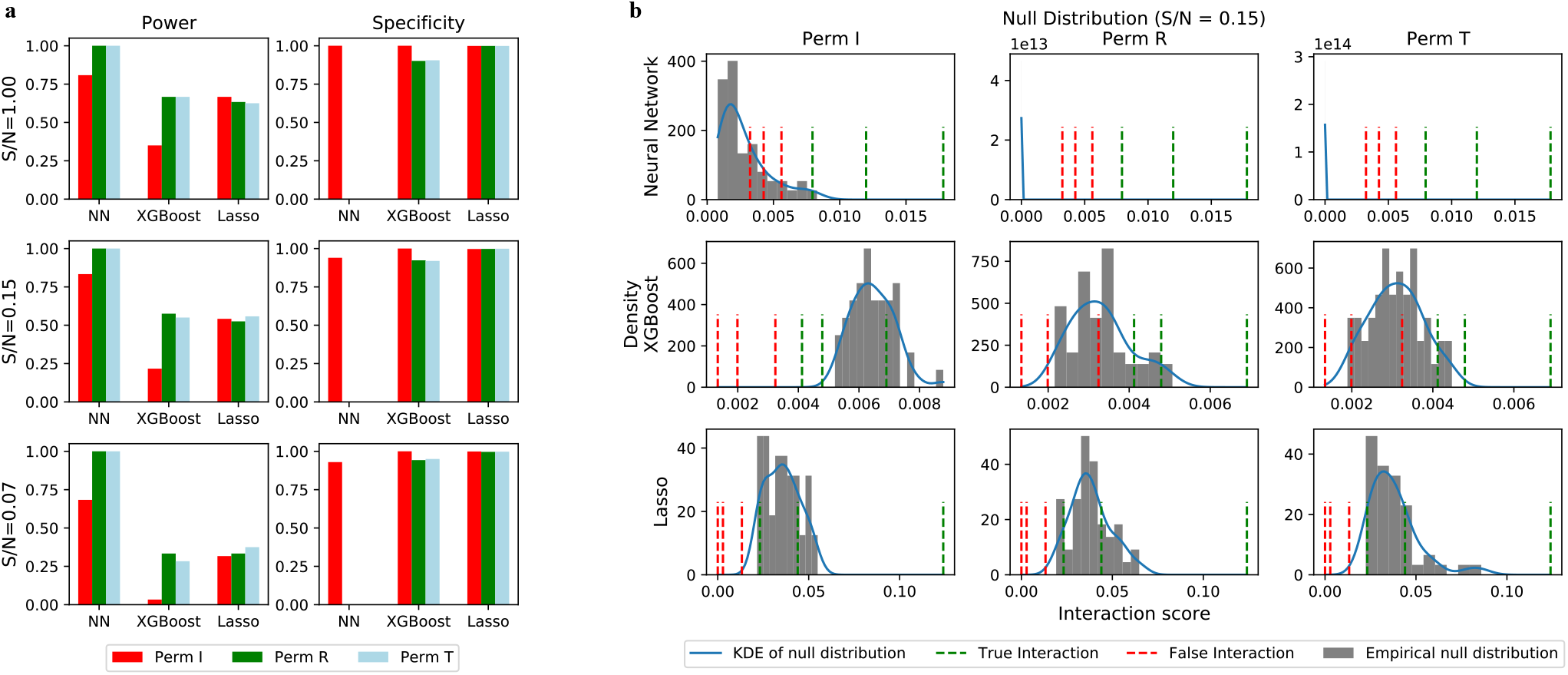
**(a)**: Comparison of the power (true positive rate) and specificity (true negative rate) of different methods on simulation datasets with complex interactions. We consider interaction as positive (i.e. detected) if its p-values is less than 0.05. NNs have greater power and similar specificity compared with Lasso and XGBoost, when interactions are tested with the proposed Perm I. With existing permutation approaches (Perm R and Perm T), NNs have specificity 0 in all settings. **(b)**: Examples of null distributions for the three different permutation methods when S/N =0.15. We visualize each null distribution with the empirical distribution and the corresponding kernel density estimate (KDE). Green/red dashed lines show three true/false interactions for illustration. For NNs (top row), the null distribution generated by Perm I correctly rejects false interactions, while Perm T and Perm R consider all false interactions positive.

**Figure 3.**
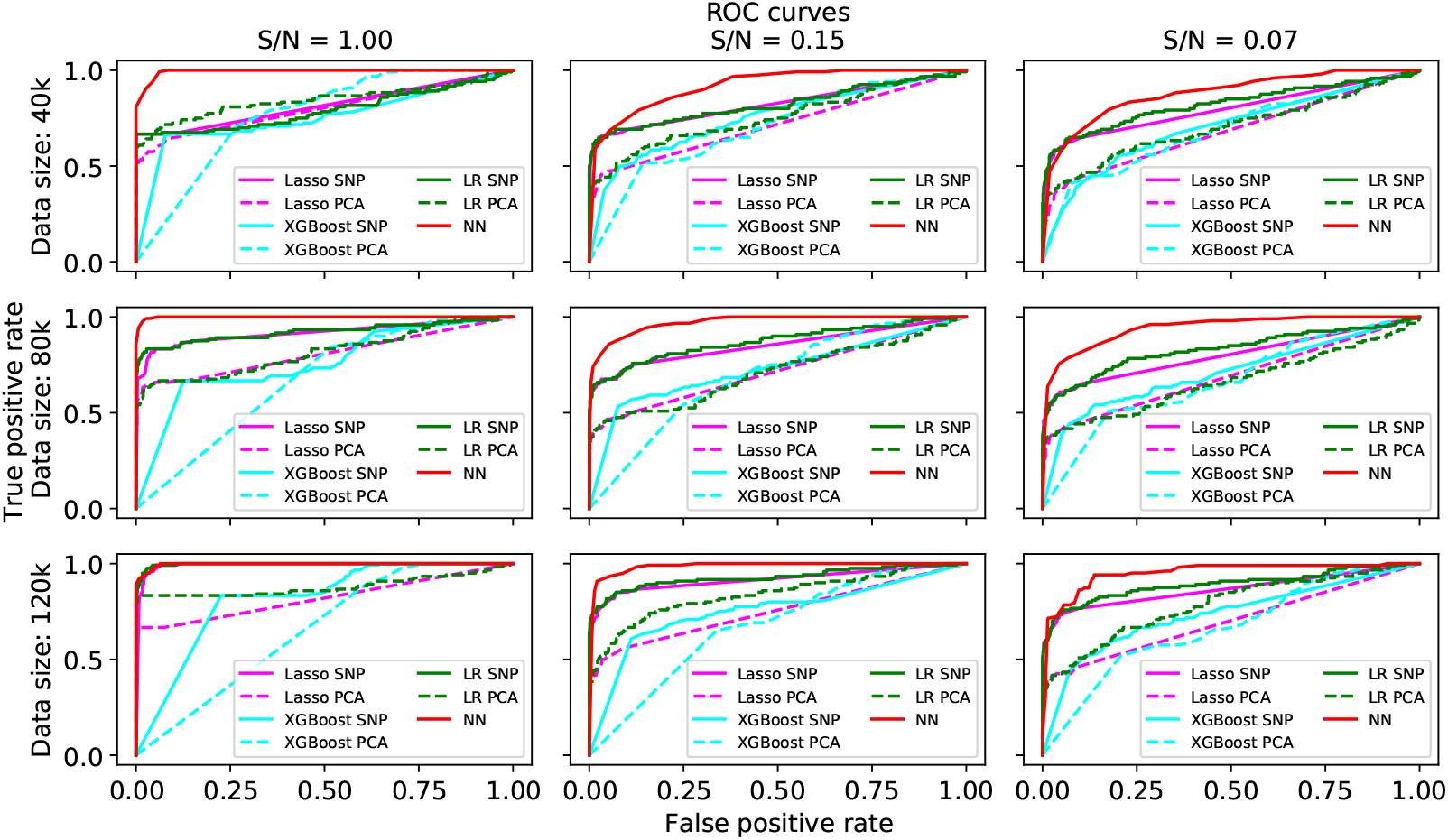
Comparison of the methods in terms of their ROC curves, as a function of dataset size (rows: 40,000, 80,000 and 120,000) and signal-to-noise ratio (S/N, columns). The NN approaches (red lines) are consistently better than existing approaches.

#### The new permutation method works well where the existing methods fail for neural networks

The proposed NN approach has a clearly greater power than the alternative methods (XGBoost and Lasso), while having roughly the same specificity when interactions are measured with the proposed permutation test (Perm I in Figure 2a). By contrast, NNs with existing permutation approaches, e.g., Perm R and Perm T in Figure 2a, have specificity 0 in all settings, as the null distributions severely underestimate the interaction scores under the null hypothesis and consider all pairwise interactions positive (top row in Figure 2b). Intuitively, interactions are estimated between hidden nodes in the gene layer, and gene representations can change dramatically after excluding main effects which usually represent most of the signal, which makes interaction detection challenging or impossible. We regard interactions as positive if their nominal p-value from permutation is less than 0.05 in Figure 2a (see results with other thresholds in the Supplementary Figure 1). In Figure 2b, we visualize example null distributions for different permutation methods when S/N =0.15 (see other S/Ns in the Supplementary Figure 2). For NNs, the null distribution of Perm I can reject false interactions correctly, by retaining the predicted main effect as part of the regression target, while Perm T and Perm R consider both true and false interactions positive (top row in Figure 2b). All permutation methods generate similar null distributions for Lasso (bottom row in Figure 2b), but Perm I overestimates interaction scores under the null hypothesis for XGBoost (middle row in Figure 2b), thus it has a lower power but a higher specificity compared with alternatives (Figure 2a).

#### NNs can detect interactions more accurately than reference methods

We observe that in all settings the NN has a larger area under the ROC curve (AUROC) than the baselines in Figure 3. This is mainly due to the fact that NNs make use of all SNPs to represent genes in a supervised manner (i.e., taking the phenotype into account), which is more informative than selecting a single SNP (e.g., top SNP) or representations learned without supervision (e.g., PCA). In addition, we see that the PCA is generally worse than the top SNP approach, which indicates the importance of using the phenotype when learning gene representations. The LR and Lasso perform better than the boosting tree when the S/N is small. Hence, we hypothesise that the interactions used in the simulator can be approximated reasonably well by the multiplicative interaction.

### 2.3 Interaction discovery in a cholesterol study

#### NNs identify 16 candidate gene-gene interactions for the HDL phenotype in the UK Biobank

We apply the NN interaction detection method on the UK Biobank dataset to demonstrate its ability to identify novel interactions. We select 17,168 SNPs (after quality control) from 65 genes which have previously been found to have large main effects for the NMR metabolomics phenotype, and we select the phenotype HDL as the dependent variable. Note that selecting genes based on their main effects does not bias the assessment of the significance of interactions using permutation. We provide the preprocessing details in Methods 4 section. We split the preprocessed data into 70% for training and 30% for testing. After we train the model on the training set, gene-gene interaction scores are estimated on the test set. We rank the interaction pairs according to their interaction scores from high to low, and we construct null distributions via the permutation test. The Q-Q plot of interaction scores on observed versus permuted datasets is shown in Figure 4a, from which we observe that the top 16 interactions significantly deviate from the expectation under the null hypothesis; we refer to these as candidate interactions. The false discovery rate of the candidate interactions set is estimated to be 0.344 (Figure 4b), which means that 9 out of 16 candidate interactions are expected to be true findings. We list all candidate interactions and their corresponding interaction scores in the first two columns of Table 1.

**Figure 4.**
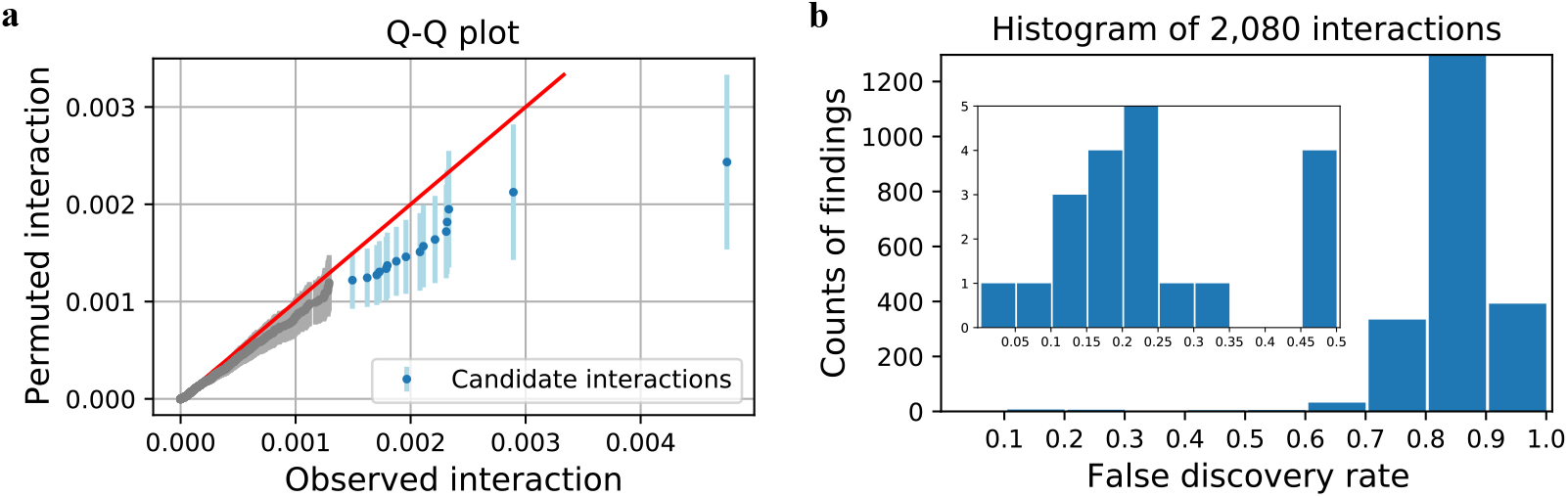
**(a)**: The Q-Q plot of Shapley interaction scores on observed vs. permuted data. The bars cover 95% of interaction scores under the null hypothesis. We observe that the top 16 interactions (blue) deviate from the null distribution. **(b)**: The histogram of interactions for different false discovery rates (FDR). The inner panel zooms in to FDR < 0.5 region. Each bar represents how many more interactions we will consider as positive if we increase FDR; e.g., 5 more findings are included by increasing FDR from 0.20 to 0.25. The top 16 interactions have FDR around 0.34, and we include these candidate interactions for a detailed analysis and replication.

**Table 1.** Detailed results for the top 16 candidate interactions identified in the UK Biobank. First column shows the corresponding pair of genes, second column the interaction score and the false discovery rate. The p-values using the top-SNP method are shown in the third column, where bold highlights p-value < 0.05. The replication results on FINRISK (p-values) are shown in the fourth column, where ‘NA’ means that the required SNPs were not available in FINRISK and the hence it was not possible to replicate the interaction. The last column gives a reference from previous literature (if any), where ‘Yes’ and ‘No’ denote whether the interaction is known/found or not, and ‘NA’ denotes that there are no related references.

| Genes | Interaction Score (FDR) | Multiplicative top-SNPs p-value | FINRISK replication p-value | Reference |
| --- | --- | --- | --- | --- |
| *ABCA1, CETP* | 4.750 (0.000) | 0.201 | **0.034** | No<sup>24</sup> |
| *LPL, LIPC* | 2.894 (0.072) | 0.257 | **0.000** | NA |
| *LPL, CETP* | 2.332 (0.218) | 0.094 | 0.800 | Yes<sup>26,29</sup> |
| *CETP, FADS1* | 2.318 (0.169) | 0.256 | 0.991 | NA |
| *LIPC, CETP* | 2.309 (0.139) | **0.012** | **0.000** | No<sup>25</sup>, Yes<sup>27</sup> |
| *CETP, PLTP* | 2.212 (0.151) | 0.987 | **0.014** | NA |
| *ABCA1, PLTP* | 2.111 (0.169) | 0.411 | 0.139 | Yes<sup>28</sup> |
| *CETP, HAL* | 2.081 (0.160) | 0.836 | NA | NA |
| *SCARB1, CETP* | 1.959 (0.195) | **0.014** | **0.006** | NA |
| *GALNT2, LPL* | 1.876 (0.218) | 0.109 | NA | NA |
| *LPL, APOA5* | 1.798 (0.239) | 0.320 | 0.915 | Yes<sup>29</sup> |
| *ABCA1, FADS1* | 1.788 (0.226) | 0.659 | 0.938 | NA |
| *LIPC, APOA5* | 1.729 (0.240) | 0.310 | **0.000** | NA |
| *CETP, GCK* | 1.706 (0.236) | 0.394 | NA | NA |
| *CETP, GCSH* | 1.623 (0.269) | 0.520 | NA | NA |
| *ABCA1, APOA5* | 1.494 (0.344) | 0.967 | 0.748 | NA |

Next we investigate if the candidate interactions detected by the NN could have been found by other methods. For the earlier method referred to as the top-SNP method, we train a bivariate LR with a multiplicative interaction term for each candidate,

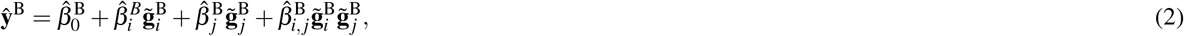

where the 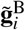 and 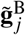 denote genes *i* and *j* represented by their top SNPs, and the superscript B refers to the UK Biobank dataset (in contrast with superscript F that will be later used for the FINRISK data). The significance of coefficient 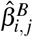 is used to check if the interaction could be detected by the top-SNP method, and the p-values are reported in the 3rd column of Table 1. We see that only 2 out of 16 candidate interactions were nominally significant using the top-SNP approach. This indicates that NNs can find interactions missed by the standard approach.

In previous studies (last column, Table 1), no interaction effect between *ABCA1*-*CETP*, and *LIPC*-*CETP* were found in a Japanese^24^ and a Chinese Han cohort^25^ respectively, where only one SNP was used to represent the gene in studies. Interaction between *LPL*-*CETP* was found in an association study on mic^e26^. Some of the interactions should exist according to the underlying biological mechanisms, such as *LIPC*-*CETP*^27^ and *ABCA1*-*PLTP*^28^, but they were not found in any association study. And finally, we found two interactions already existing in KEGG^29^ genome-encyclopedia, first being *LPL*-*CETP* in the glycerolipid metabolism pathway, and second being *LPL*-*APOA5* in the PPAR signaling pathway.

#### Six of the candidate interactions replicate in the FINRISK dataset

We use an independent dataset, FINRISK, to verify if the 16 candidate interactions detected in the UK Biobank are replicable. We select the same SNPs from the FINRISK, and calculate gene representations according to the model learned with the UK Biobank. For replication, we first calculate the residual in the FINRISK by subtracting the main effects from the phenotype value for a given pair of genes, i.e., 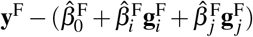. Then, we estimate the interaction pattern for the gene pair in the UK Biobank by training two NNs: a fully connected MLP, 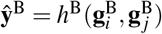 and another NN that only contains non-linear main effects without any interactions, 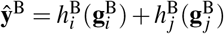. We represent the interaction learned from the UK Biobank by the difference between these two NNs, 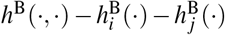, i.e., this is the part that cannot be explained by main effects. Finally, we use the following linear regression to test if the interaction pattern learned from the UK Biobank can explain the observed residual in the FINRISK:

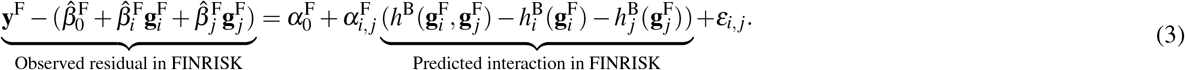

If 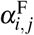 is significantly greater than zero, we regard the interaction between genes *i* and *j* as replicated. The replication p-values for each interaction are shown in Table 1. The set of SNPs involved in the interaction is available in the FINRISK for only 12 out of the 16 candidate interactions, and therefore it is possible to try to replicate only these 12 interactions. Out of the 12 possible interactions, six finally replicate significantly in the FINRISK, with a FDR around 0.068.

In Figure 5, we visualize 3 replicated interactions and the corresponding fits for the linear regressions in Eq. 3 (see the remaining 3 interactions in the Supplementary Figure 5). From the linear regression (first row, Figure 5), we notice that the interaction pattern learned from the UK Biobank can explain the observed residual in the FINRISK. We use blue, purple, and yellow background colorings to represent individuals with low, medium, and high interaction values respectively, and mark those individuals in the visualization of the learned interaction pattern with the same color. From the visualizations (Bottom row, Figure 5), we notice that the shape of the low interaction regions, the pink, can be elliptical, concave, and even multi-modal, which can not be approximated well by a multiplicative interaction. For example, the interaction effect between *LIPC* and *LPL*is negative only if the representation of *LIPC* is <-1.5 and at the same time LPL is in the range (0.0,1.0), and the interaction between *CETP* and *SCARB1* has a complex multi-modal shape. We further notice that even if these interactions are replicated significantly, the effect sizes are relatively small, increasing *R*^2^ on average by 0.7% (ranging from 0.3% to 3.5%) in the test set and by 1.7% (ranging from 0.7% to 5.5%) on the whole dataset (see details in Supplementary Table 1).

**Figure 5.**
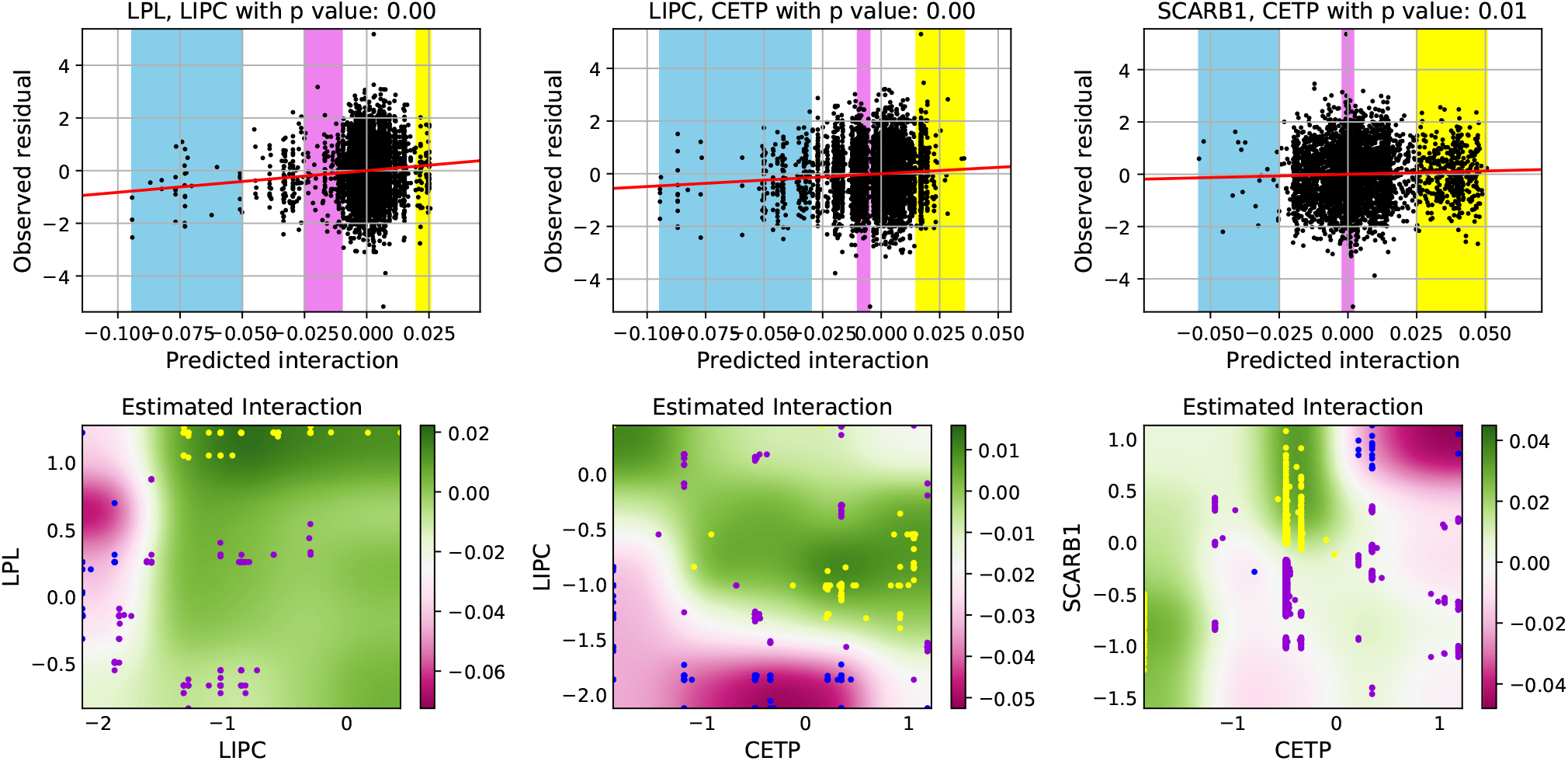
Visualization of three interactions replicated in the FINRISK dataset. **Top**: X-axis represents the interaction predicted in FINRISK using the model trained on the UK Biobank. Y-axis shows the residual after removing the main effects in the FINRISK dataset, see Eq. 3. The interactions learned from the UK Biobank (x-axis) significantly explain the variation in the phenotype of FINRISK that remains after excluding the main effects (y-axis). **Bottom**: Heatmaps representing the estimated interaction (background color) between two gene representations (x-axis and y-axis) learned from the UK Biobank. Blue, purple, and yellow backgrounds in the top panels represent individuals with low, medium, and high predicted interaction values, and they are shown with dots in the bottom panels. It is clear that the replicable interactions have complex shapes which can not be approximated well by multiplicative interactions.

## 3 Discussion

Existing approaches for detecting interactions in a GWAS do not reveal all gene-gene interactions because 1) genes are typically represented by the single most significant SNP, and 2) only restricted forms of interaction (e.g. multiplicative) are considered. Therefore, the statistical power can be limited for complex gene-gene interactions and important interactions may be ignored. Here, we resolved these shortcomings with a novel deep learning approach. NNs are generally well-known for their ability to learn high level representations from low level features with suitable architectures that reflect prior knowledge of the application domain. Furthermore, as a universal function approximator, a NN can model any flexible relation between genes and phenotypes, including interactions. Despite these benefits, NNs have not been widely used to detect interactions in GWAS because it is neither trivial to uncover the interactions from a NN nor assess the significance of the findings. We defined a gene interaction NN using a structured sparse architecture, which learned a representation of each gene from all the corresponding SNPs, instead of considering a designated top SNP or ignoring information in the phenotype (as, e.g., PCA, does). From this network, by applying a mathematically principled interpretation method, Shapley interaction scores, we were able to estimate the flexible gene-gene interactions as a post-hoc step. In addition, we designed a new permutation procedure to assess the statistical significance of interactions, which is essential to control the false discovery rate of findings.

The proposed deep learning framework has many attractive properties, but also some shortcomings. First, although the permutation distribution of gene-gene interaction scores gives decent results in practice, whether the permutation distribution is well-calibrated still requires further theoretical investigation. Second, the framework could easily be extended to detect higher-order interactions by calculating higher-order Shapley interaction scores, but this would require *O*(*M^t^*) forward passes for *t*-order interactions among *M* genes, and therefore computation could be a bottleneck. We noticed that although NNs can detect novel interactions, the estimated effect sizes are relatively small. This could be improved by improving the architecture of the NN or using more informative priors. Lastly, we point out that although NNs have the highest power among various baselines on simulated datasets with complex interactions, the power of NNs is smaller than pairwise top-SNP approaches on the UK Biobank (Supplementary Figure 4). We hypothesise that most interactions in the real-world data can be approximated well by multiplication between top SNPs. Therefore NN should not used as a substitute for the default top-SNP approach, but rather as a complementary tool to find interactions that would otherwise be missed. Indeed, with real world genetic data, we showed that our NN Framework can find interactions that cannot be detected via ordinary SNP based approaches, for example, 4 of the 6 replicable interactions could not be detected by a linear regression with top-SNP representations (Figure 5).

## 4 Methods

### Data preprocessing

UK Biobank (UKB) is a prospective cohort of approximately 500,000 individuals from the United Kingdom, which consists of rich genotype and phenotype information of each individual^15^. We select 424,389 individuals that pass sample quality control with 65 genes which have previously been found to have high main effects on cholesterol phenotypes such as the HDL^23,30^, and 10 out of 65 genes are used on simulated datasets. We select all SNPs included within the 65 chosen genes with 10kbp flanking regions added to both ends of each gene. Quality control is conducted by filtering out SNPs with missing rate > 0.05, MAF < 0.01, pHWE < 10^−6^ and info score (imputation quality) < 0.4, leaving 17,168 SNPs. To reduce collinearity, we prune SNPs to keep 90% of total variance according to the linkage disequilibrium (LD) score^31^, finally amounting to 4,322 SNPs. We use HDL as the target phenotype to be associated with the selected genes. We regress out age and gender covariates as well as the first 10 genotypic principal components to remove population structure. The residuals are quantile normalized before using them as the dependent variable.

To replicate the positive gene-gene interactions detected from the UK Biobank, we select an independent dataset from the FINRISK project (DILGOM07 subset) which includes a cohort of unrelated individuals aged 30-59 years participating in a study on coronary heart diseases in Finland^16^. We select 4,620 individuals that pass the sample quality control, and select the same set of SNPs as in UK Biobank for each gene in order to reuse the NN gene representations learned on the UK Biobank dataset. We ignore genes that do not contain the SNPs used in the UK Biobank experiments, which reduces the number of candidate interactions for replication to 12 (Table 1). We choose HDL as a regression target, and apply the same preprocessing steps on HDL as we did on the UK Biobank.

### Notation

We assume that there are *N* individuals and *M* genes with *m_i_* SNPs for gene *i*, and 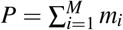 SNPs in total. We denote the target variable, i.e., one chosen phenotype, by a vector **y** = (*y*_1_,…, *y_N_*)^*T*^ and the covariate matrix containing all SNPs of the selected genes by 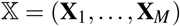, where **X**_*i*_ = (**x**_*i*1_,…, **x**_*im_i_*_) represents the matrix of SNPs of gene *i* and *x_ij_* = (*x_ij_1*,…, *x_ijN_*)^*T*^ is an *N*-dimensional column vector that represents the *j*th SNP of gene *i* for all individuals. We use *X_k_* to represent a *P*-dimensional row vector of all SNPs of individual *k*, such that 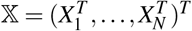.

### Gene interaction NNs

The architecture of a NN reflects inductive biases of the model, which is one of the keys to the success of NNs. For example, the convolutional and pooling layers designed for computer vision learn functions that are translation invariant^32^. In genetics, the effect of SNPs on a phenotype can be abstracted into a two-stage procedure: 1. the SNPs within a gene jointly affect the gene expression; 2. the expressions of multiple genes affect phenotypes eventually. We propose a novel NN architecture with structured sparsity that leverages these two mechanisms. Mathematically, a structured sparse neural network model can be written as

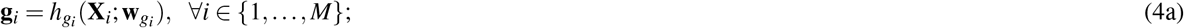

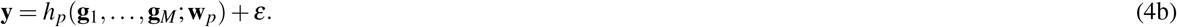

where *h_gi_*(·) represents a multilayer perceptron (MLP) parametrized by **w**_*gi*_ and the output **g**_*i*_ is an *N*-dimensional vector that learns the representation of gene *i* from its SNPs **X**_*i*_. *h_p_*(·) is another MLP that models the relations between genes and the phenotype **y**, and *ε* represents the part of the phenotype that cannot be explained by genetic information. We demonstrate the architecture in Figure 1, where the hidden layer that contains gene representations **g** is called the gene layer. The architecture is structured sparse because SNPs from different genes are disconnected until the gene layer. All MLPs in Eq. 4 are trained in an end-to-end manner by reformulating the model as

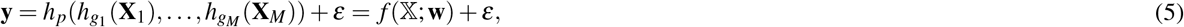

where **w** represents all parameters in the model. Eq. 5 only takes the SNPs 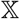 and phenotype **y** as inputs during training. Empirically, we find that using small MLPs for both *h_gi_*(·) and *h_p_*(·) usually has a better accuracy than large architectures. We use one hidden layer MLP with architecture *m_i_* – 10 – 1 (i.e., *m_i_* nodes in the input layer, 10 nodes in the hidden layer, and 1 node in the output layer) for *h_gi_*(·) and 10 – 100 – 1 for *h_p_*(·) in simulation datasets, and use *m_i_* – 10 – 1 for *h_gi_*(·) and 65 – 200 – 1 for *h_p_*(·) in the UK Biobank dataset, where 65 is the number of genes considered.

### Bayesian inference of NNs

Different from deterministic NNs, Bayesian NNs^33^ are defined by placing a prior distribution *p*(**w**) over the weights, and instead of finding a point estimate of the weights by minimizing a cost function, a posterior distribution of the weights is calculated conditionally on the data. Here we adopt the Bayesian approach because it makes inference more robust by averaging over multiple parameter values instead of using a point estimate and allows for incorporating prior knowledge about sparsity and effect size which helps when learning the weak genetic signals. The Bayesian approach could also be used to quantify uncertainty and assess significance in principle, but the calibration and use of posterior distributions to address the multiple testing problem are still active research topics; hence, we instead use permutation testing to assess significance here because that is a well-established method in the context of GWAS.

Given a dataset 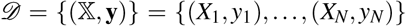 and likelihood function 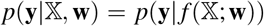, learning Bayesian NN means computing the posterior distribution 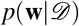 according to the Bayes’ rule

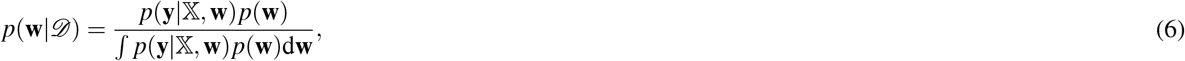

which is analytically intractable because of the integration in the denominator. Variational inference can be used to approximate the true posterior 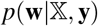 with a simpler distribution, *q_ϕ_*(**w**), by minimizing the Kullback–Leibler divergence 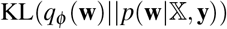. This is equivalent to maximizing theELBO (Evidence Lower BOund)

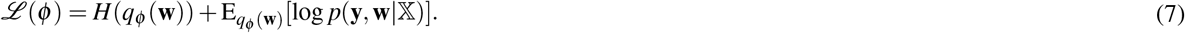

The first term in Eq. 7 is the entropy of the approximated posterior, which can be calculated analytically for many choices of *q_ϕ_*(**w**). The expected log likelihood term is intractable, but can be estimated by re-parametrising^34^ the approximated posterior *q_ϕ_*(**w**) by a deterministic and differentiable function **w** = *g*(*ε; ϕ*) with *ε* ~ *p*(*ε*), such that *E*_*qϕ*(**w**)_[log *p*(**y, w|X**)] = E_*p*(*ε*)_[log *p*(**y**, *g*(*ε; ϕ*)|**X**)]. Then we replace the second term with its stochastic estimator, such that

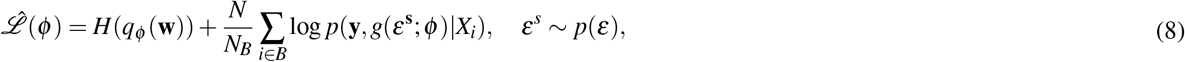

which can be optimized by stochastic gradient descent with batch size *N_B_*.

We use an independent spike-and-slab prior for each weight with Automatic Relevance Determination (ARD)^35^, which means that all of the outgoing weights 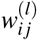 from the same node *i* in layer *l* share the same scale 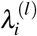

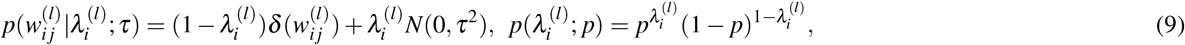

where 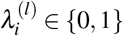, and *δ*(·) is the Dirac delta function. Spike-and-slab prior with ARD induces sparsity w.r.t. the input features (SNPs), which is especially beneficial in our case because the set of all SNPs for each gene can be large. Two hyper-parameters in the spike-and-slab prior are the sparsity level *p* and the global scale *τ*. Moreover, given a sparsity level *p*, the global scale *τ* is the only factor that affects the proportion of variance explained (*R*^2^) of the model, which is easier to interpret than global scale *τ*^36^. Thus hyper-parameters *p* and *τ* are chosen by a 5-fold cross-validation on *p* and *R*^2^ on the training set. The approximate posterior 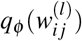 is also an independent spike-and-slab distribution, such that

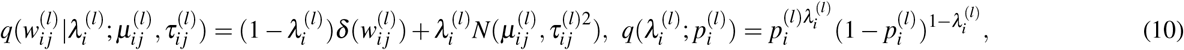

where the variational parameter *ϕ* contains the mean 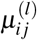 and standard deviation 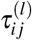 of the Gaussian distribution for each weight and the slab probability 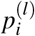 for each node.

For discrete random variables, such as the Bernoulli distribution used here, the reparametrization trick cannot be applied directly. Thus a continuous approximation of discrete Bernoulli, the concrete Bernoulli distribution^37^, is used instead. Sampling from the concrete Bernoulli is equivalent to using a deterministic function of the Bernoulli parameter *p*, and random noise *ε* from the Logistic distribution

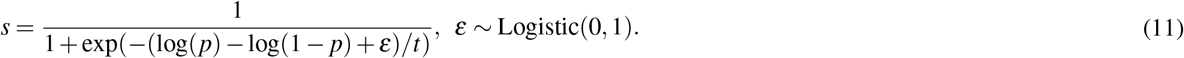

Here *t* is the temperature which controls the trade-off between the approximation quality and differentiability. Decreasing *t* improves the accuracy of the approximation, at the cost of having sparser gradients. We use the concrete Bernoulli with *t* = 0.1 to approximate the posterior of discrete local scale parameters, i.e., 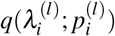, in experiments.

### Deep ensemble of NNs

The loss surface (Eq. 8) of NNs is highly multi-modal, and the stochastic variational inference used to learn the Bayesian NN’s posterior can only find one of the modes^17^. An easy way to capture multiple modes and thus improve the approximated posterior is through a deep ensemble^38^, which repeats the same training procedure multiple times with different initialization of parameters and then averages different NNs during testing. We apply this trick for all experiments, where we use an ensemble of 30 NNs trained with different initialization. Deep ensemble is computationally inefficient, but it can be parallelized easily.

### Gene-gene interaction score

Shapley interaction score^10^ is a well-axiomatized interaction score for input features motivated by the Shapley values in game theory. We denote the set of all features by *F*, a feature *i* ∈ *F*, and a feature set *S* ⊆ *F*. We define the interaction effect between feature *i* and *j*, with feature set *S*, of a neural network *f* at a data point *X_k_* to be

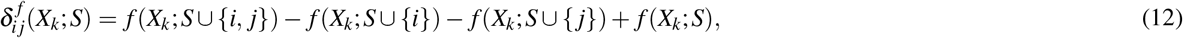

where *f*(*X_k_*; *S*) is the prediction at *X_k_* when only features in *S* are used, which often requires retraining the NN multiple times. A common approximation is to replace the absent features (i.e., *F*\*S*) by the corresponding values in a baseline *C_F\S_*, such that

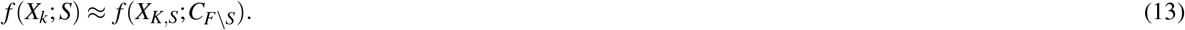

The baseline is usually set as the empirical mean of each feature, which is used in this work as well, but more informative baselines can also be applied. The Shapley interaction score 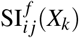 is the expectation of 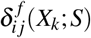,

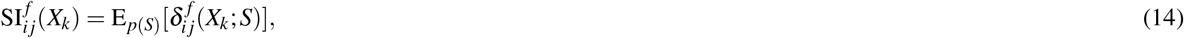

over a uniformly random chosen feature set *S* from *F*. This could be computationally expensive when the dimension |*F*| is high. We use a Monte-Carlo procedure^39^ to approximate 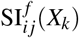 by a small number of samples of *S*.

Shapley interaction score 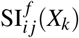 is a *local* interaction effect measure, representing the interaction effect between feature *i* and *j* of function *f*(·) at data point *X_k_*. To aggregate the *local* interaction effect at different data points into a *global* interaction effect, i.e., shared by the whole data domain, we use take the expectation of 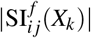 w.r.t. the empirical data distribution *p*(*X*), such that

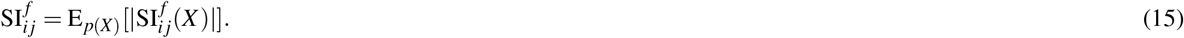

Instead of the interaction between individual input features (SNPs), we are interested in interactions between genes, which are represented by the hidden nodes of the gene layer. Thus the interaction between genes *i* and *j* can be calculated by the Shapley interaction score between between **g**_*i*_ and **g**_*j*_ in *h_p_*(**g**_1_,…, **g**_*M*_; **w**_*g*_) on Eq. 4b, where the gene representations, **g**, are calculated in lower layers on Eq. 4a. For a deep ensemble of Bayesian NNs, instead of having a point estimator of function *f*(·; **w**), we have a posterior distribution of functions *q*(*f*) induced by the posterior of the weight *q*(**w**). Thus the interaction score of Bayesian NN is the expectation of 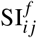 w.r.t. *q*(*f*), which we estimate by taking the average of *N_f_* samples drawn from the posterior:

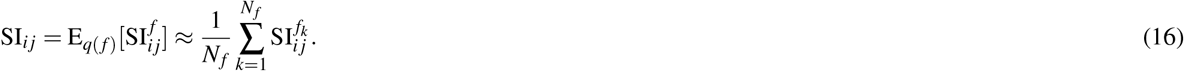

In experiments, we draw 10 NNs from the posterior of each Bayesian NN, and thus we have in total 300 NNs in the whole ensemble (i.e., 30 BNNs times 10), to estimate the interaction scores between genes.

### Permutation test of interaction

The alternative hypothesis that we want to verify is that interaction effects between genes are non-zero. Thus the corresponding null hypothesis is: genes only have main effects on the phenotype without any interactions. We use a structured sparse linear regression (shown in Figure 1) to represent the null hypothesis model (i.e., main effects of genes):

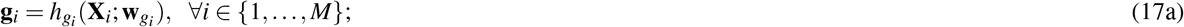

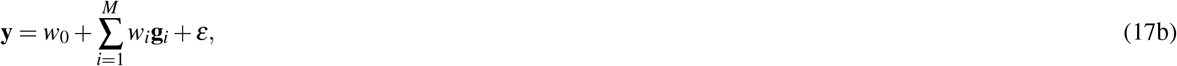

where the NN *h_p_*(·) in Eq. 4b is replaced by a linear regression. When *h_gi_*(·) is reduced to a linear regression, the null hypothesis (Eq. 17) will be a reduced rank regression^23^ with rank *M*.

We propose a new permutation method to generate permutation datasets for NN training. We first permute the residual of the null hypothesis (Eq. 17), which removes any interaction effects from the data. Then we define the dependent variable as the sum of the predicted phenotype of Eq. 17 and the permuted residual, which retains the main effect. We do not permute the independent variable. The permutation procedure (**Perm I**) can be summarized as follows:

1. Estimate residuals, **r** = **y** – **ŷ**, by fitting the null hypothesis model in Eq. 17;
2. Permute the interaction effect of data by permuting **r** to obtain **r**^*π*^;
3. Obtain the permutation target by **y**^*π*^ = **ŷ** + **r**^*π*^;
4. Compute Shapley interaction score of NNs (Eq. 4) trained on permutation datasets 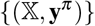;
5. Repeat Steps 2-4 several times to obtain an empirical null distribution.

We compare against two existing permutation methods on simulated data:

**Perm T**: use permuted dependent variable as target^11^:

1. Permute the dependent variable **y** to obtain **y**^*π*^;
2. Compute Shapley interaction score of NNs (Eq. 4) trained on permutation datasets 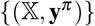;
3. Repeat Steps 1-2 several times to obtain an empirical null distribution.

**Perm R**: use permuted residual as target^14^:

1. Estimate residuals, **r** = **y** – **ŷ**, by fitting the null hypothesis model in Eq. 17;
2. Permute the interaction effect of data by permuting **r** to obtain **r**^*π*^;
3. Obtain the permutation target by **y**^*π*^ = **r**^*π*^;
4. Compute Shapley interaction score of NNs (Eq. 4) trained on permutation datasets 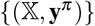;
5. Repeat Steps 2-4 several times to obtain an empirical null distribution.

Perm T removes the interaction as well as the main effect. Perm R only permutes the interaction effect, however it also fails to keep the main effects intact in the permuted datasets.

We incorporate two approaches for multiple testing correction in the permutation procedure: Max-T method^20^ and false discovery rate (FDR) control^21^, in simulated datasets and real-world applications, respectively. For Max-T method, we collect the maximal Shapley interaction score across all interaction scores for each permutation in Step 4. This provides a single empirical null distribution for all interaction pairs to conduct hypothesis tests. For FDR, we record the Shapley interaction scores of all interaction pairs from high to low for each permutation in Step 4, which allows us to calculate the FDR given a decision threshold of interaction score. FDR has less stringent control of type I error but greater statistical power than the Max-T method. Permutation testing is computationally heavy as it requires training multiple NNs. However, like a deep ensemble, permutations are easy to parallelize by applying different random seeds to generate local permutation targets.

## Supporting information

Supplementary and Additional Results

## Data availability

UK Biobank data are available to registered investigators under approved applications [http://www.ukbiobank.ac.uk]. A permission to use the FINRISK data can be applied from THL (Finnish Institute for Health and Welfare). Other relevant data are available from the corresponding author upon request.

## Code availability

The source code is available at [https://github.com/tycui/GWAS_NN].

## Acknowledgements

The work used computer resources of the Aalto University School of Science Science-IT project. This work was supported by the Academy of Finland (Flagship programme: Finnish Center for Artificial Intelligence, FCAI, and grants 319264, 292334, 286607, 294015, 336033, 321356), and by the EU Horizon 2020 (grant no. 101016775).

## Author contributions

PM and SK designed the study. TC, KM, JR, PM and SK developed the method. TC implemented the method and ran all the experiments. KM and AH preprocessed the datasets. AH, PM and SK supervised the work. TC wrote the article with contributions from all authors.

