## Supplementary and Additional Results for "Gene-Gene Interaction Detection with Deep Learning"

### Supplementary for "Gene-Gene Interaction Detection with Deep Learning"

#### Contents

**1 Additional experiments on simulated data** **2**

**2 Additional experiments on UK Biobank** **5**

**3 Visualization of replicable interactions** **7**

### 1 Additional experiments on simulated data

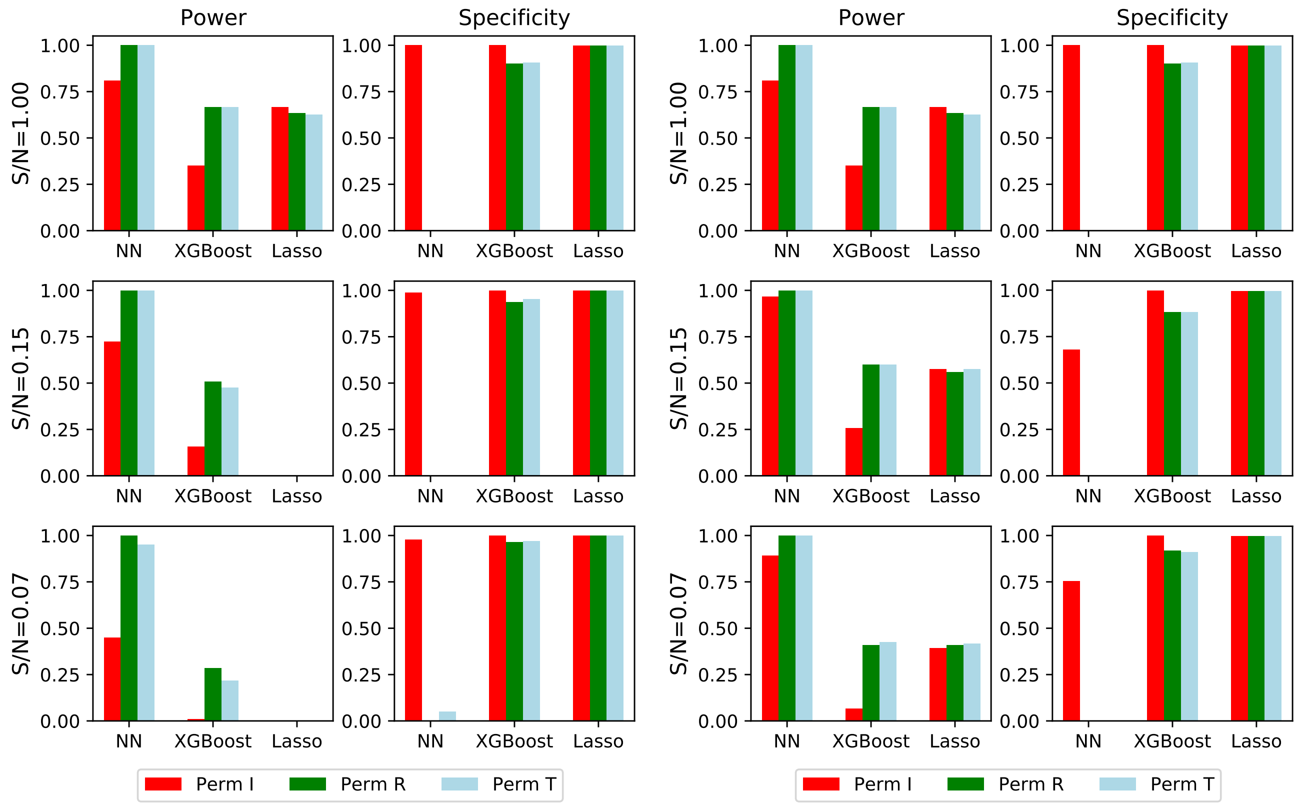


Figure 1: Comparison of the power (true positive rate) and specificity (true negative rate) of different methods. We consider interactions to be positive if corresponding p-values are less than 0.02 (left) and 0.1 (right).

In Figure 1, we compare the power and specificity of different methods on simulated datasets. We regard interactions are positive if their nominal p-value from permutation is less than 0.02 and 0.1. We notice that by increasing the p-value threshold from 0.02 to 0.1, the power of each method (especially for low S/Ns, e.g., 0.15 and 0.07) increases, while the corresponding specificity decreases. For NNs, Perm T and Perm R have specificity 0 and power 1 in most settings, as they severely underestimate the interaction scores under the null hypothesis and consider all pairwise interactions positive. We observe that although NNs have a specificity similar to or slightly smaller than the other methods, the power of detecting complex interactions is much larger.

In Figure 2, We visualize example null distributions of interaction scores for different permutation methods when S/N = 0.07 and S/N = 1.0, where we noticed that existing permutation methods underestimate the interaction scores in null distribution seriously.

In Figure 3, we visualize the ROC of two additional S/N (0.5, 0.1), where we can conclude that NN has a larger AUROC than the baselines.

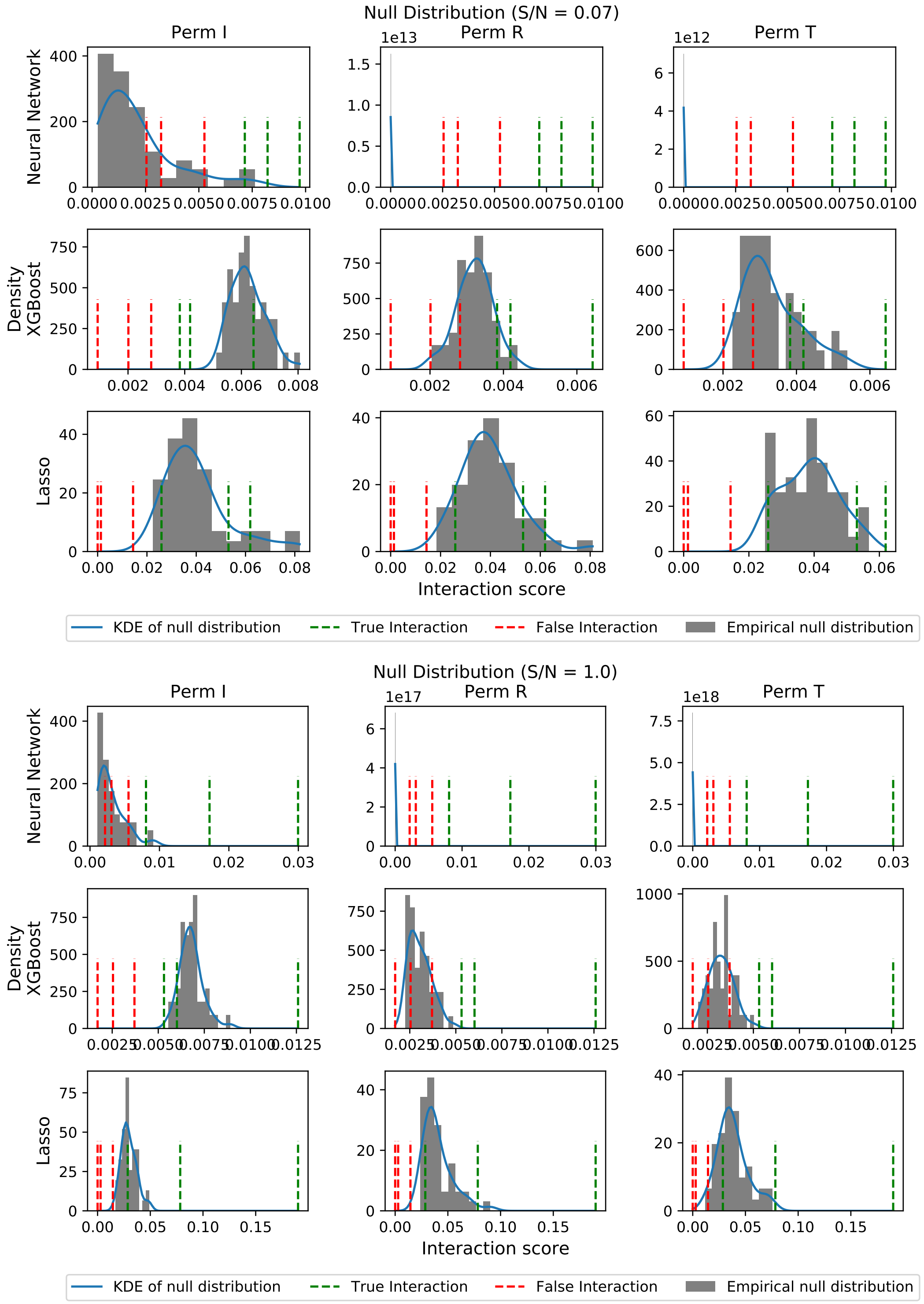


Figure 2: Visualization of null distributions from different permutation methods. We select three true interactions (green dash) and three false interactions (red dash) for illustration. We consider two signal-to-noise ratio levels: 0.07 (top) and 1.0 (bottom).

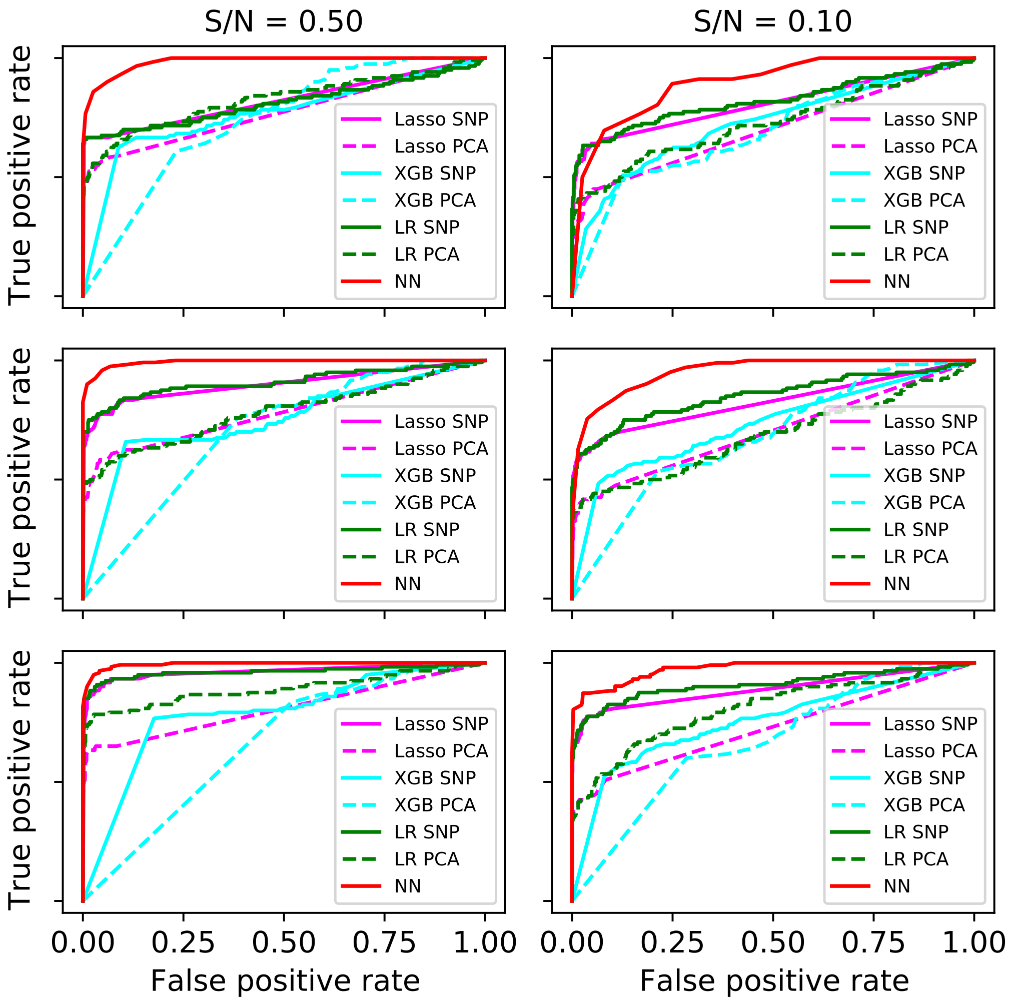


Figure 3: Comparison of ROC curves of different methods on different datasets. Different rows represent different sizes of data: 40,000, 80,000 and 120,000. Different columns represent different signal-to-noise ratios. The NN approaches (red lines) are significantly better than existing approaches.

### 2 Additional experiments on UK Biobank

Table 1: Proportion of variance explained ($R^2$) by linear regressions and neural networks on each pair of interactions. We observe that although NNs have a higher $R^2$ than LR in general, the sizes of increases are generally small.

| Genes | Whole data LR, $R^2$ (%) | Whole data NN, $R^2$ (%) | Whole data Increases (%) | Test data LR, $R^2$ (%) | Test data NN, $R^2$ (%) | Test data Increases (%) |
|---|---|---|---|---|---|---|
| ABCA1, CETP | 4.742 (0.000) | 4.774 (0.003) | 0.679 (0.067) | 4.610 (0.003) | 4.621 (0.006) | 0.238 (0.151) |
| LPL, LIPC | 2.408 (0.001) | 2.433 (0.003) | 1.018 (0.133) | 2.467 (0.002) | 2.475 (0.006) | 0.332 (0.248) |
| LPL, CETP | 5.252 (0.001) | 5.282 (0.003) | 0.583 (0.063) | 5.228 (0.003) | 5.243 (0.004) | 0.283 (0.103) |
| CETP, FADS1 | 4.173 (0.000) | 4.203 (0.003) | 0.718 (0.061) | 4.092 (0.003) | 4.109 (0.005) | 0.419 (0.123) |
| LIPC, CETP | 5.233 (0.001) | 5.281 (0.004) | 0.915 (0.085) | 5.148 (0.003) | 5.176 (0.006) | 0.535 (0.123) |
| CETP, PLTP | 4.200 (0.001) | 4.227 (0.003) | 0.648 (0.069) | 4.103 (0.003) | 4.116 (0.006) | 0.315 (0.168) |
| ABCA1, PLTP | 0.891 (0.000) | 0.898 (0.002) | 0.841 (0.227) | 0.836 (0.001) | 0.836 (0.004) | 0.001 (0.420) |
| CETP, HAL | 4.037 (0.001) | 4.066 (0.002) | 0.707 (0.059) | 3.945 (0.003) | 3.957 (0.006) | 0.324 (0.169) |
| SCARB1, CETP | 4.352 (0.001) | 4.379 (0.003) | 0.619 (0.064) | 4.193 (0.004) | 4.206 (0.007) | 0.314 (0.198) |
| GALNT2, LPL | 1.554 (0.050) | 1.582 (0.002) | 1.956 (0.362) | 1.571 (0.024) | 1.575 (0.006) | 0.275 (0.164) |
| LPL, APOA5 | 1.550 (0.102) | 1.611 (0.005) | 4.456 (0.831) | 1.567 (0.060) | 1.608 (0.006) | 2.801 (0.450) |
| ABCA1, FADS1 | 0.857 (0.000) | 0.863 (0.002) | 0.642 (0.270) | 0.825 (0.001) | 0.823 (0.004) | -0.227 (0.568) |
| LIPC, APOA5 | 1.513 (0.125) | 1.589 (0.005) | 6.299 (1.566) | 1.501 (0.063) | 1.553 (0.009) | 3.678 (0.498) |
| CETP, GCK | 4.040 (0.001) | 4.069 (0.003) | 0.729 (0.067) | 3.954 (0.003) | 3.973 (0.005) | 0.466 (0.149) |
| CETP, GCSH | 4.030 (0.001) | 4.058 (0.003) | 0.706 (0.080) | 3.939 (0.004) | 3.945 (0.006) | 0.149 (0.156) |
| ABCA1, APOA5 | 1.040 (0.081) | 1.089 (0.004) | 5.431 (0.991) | 0.991 (0.044) | 1.014 (0.007) | 2.524 (0.526) |

In Table 1, we compute the proportional of variance explained ($R^2$) of linear regression models (without any interactions) and neural networks on the whole dataset and on a separate test dataset respectively. We notice that although using neural networks can improve the variance explained, the sizes of increases are generally small. This indicates that there is room to improve by using more domain specific neural network architectures and incorporate domain knowledge via informative priors.

In Figure 4, we train a bivariate LR with a multiplicative interaction term for each interaction pair, where the gene is represented by the corresponding top SNP. We collect the negative log p-value (y axis) of the multiplicative interaction term of linear regression and the corresponding gene-gene interaction score (x axis) ofrom NNs for each interaction terms. We apply two FDR (false discovery rate) thresholds: 0.1 and 0.34. We observe that NNs can find very different interactions from the top-SNP approach, and for a certainty threshold, the number of findings from NNs is smaller than top-SNP approach. This indicate that most interactions in real-world applications (e.g., UK Biobank dataset) can be approximated well by multiplications between top SNPs, and then that is the best thing to do. Therefore the NN approach should not be used as a substitute for the default top-SNP approach, but rather as a complementary tool to find interactions that would be missed otherwise.

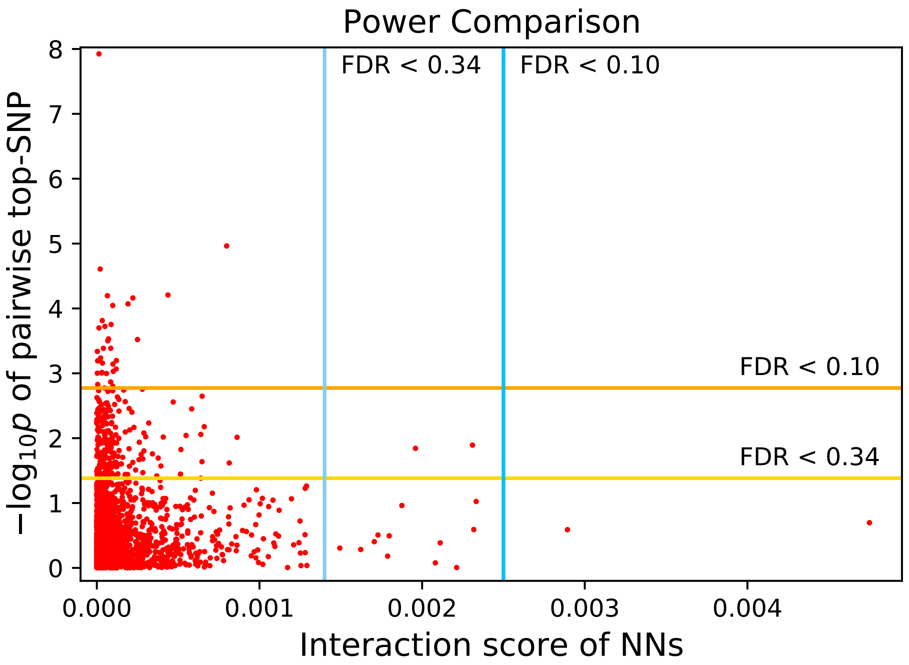


Figure 4: Compare the findings of top-SNP approach and NNs approach with same FDR thresholds. Each red dot represents one interaction pair, and the corresponding x and y coordinates are the interaction score from NNs and the negative log p-value of top-SNP regression respectively. We notice that NNs can find very different interactions from the default top-SNP regression approach, and the number of findings are smaller than top-SNP approach with the same FDR threshold.

### 3 Visualization of replicable interactions

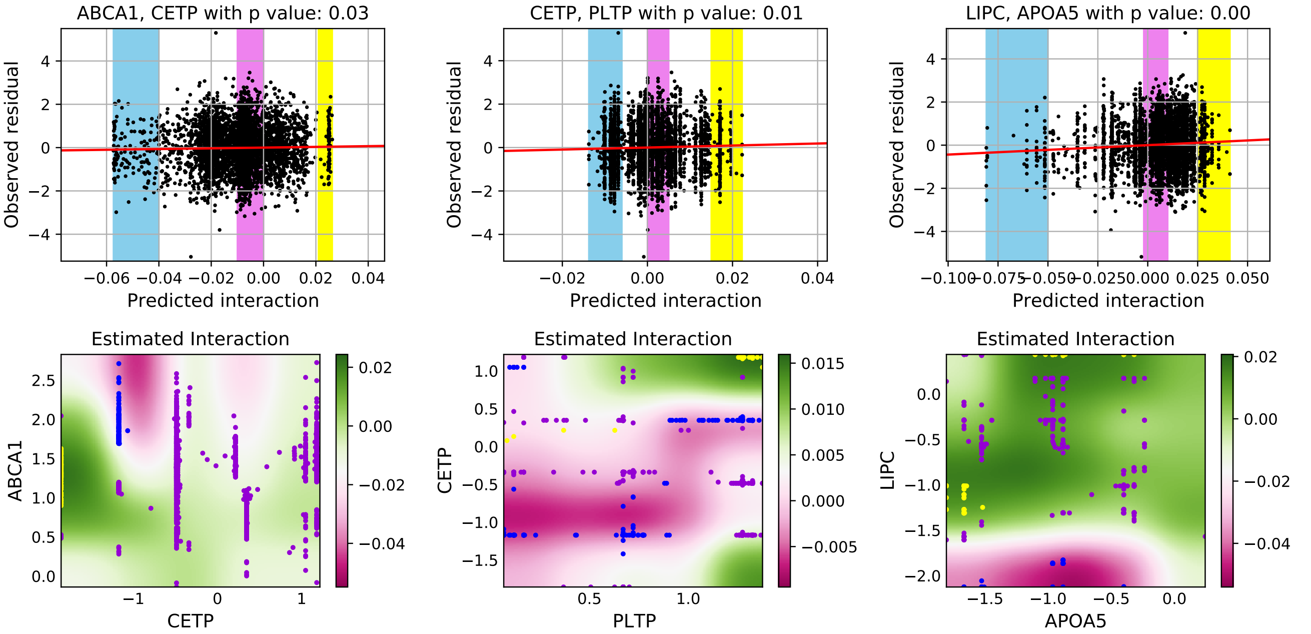


Figure 5: Replicate interactions detected from UK Biobank on FINRISK dataset. **Top**: Visualization of 3 linear regressions for replication. **Bottom**: Visualization of corresponding gene-gene interactions, where the heatmap represent the learned interaction function. Blue, purple, and yellow background colorings in top panels represent the individuals with the low, medium, and high interaction values respectively, and we mark those individuals in bottom panels accordingly.

We visualize another 3 replicable interactions and the corresponding fits for the linear regressions in Figure 5.
